# Cytokinin-CLAVATA crosstalk is an ancient mechanism regulating shoot meristem homeostasis in land plants

**DOI:** 10.1101/2021.08.03.454935

**Authors:** Joseph Cammarata, Christopher Morales Farfan, Michael Scanlon, Adrienne HK Roeder

## Abstract

Plant shoots grow from stem cells within Shoot Apical Meristems (SAMs), which produce lateral organs while maintaining the stem cell pool. In the model flowering plant Arabidopsis, the CLAVATA (CLV) pathway functions antagonistically with cytokinin signaling to control the size of the multicellular SAM via negative regulation of the stem cell organizer WUSCHEL (WUS). Although comprising just a single cell, the SAM of the model moss *Physcomitrium patens* (formerly *Physcomitrella*) performs equivalent functions during stem cell maintenance and organogenesis, despite the absence of WUS-mediated stem cell organization. Our previous work showed that the stem cell-delimiting function of the CLV pathway receptors CLAVATA1 (CLV1) and RECEPTOR-LIKE PROTEIN KINASE2 (RPK2) is conserved in the moss *P. patens*. Here, we use *P. patens* to assess whether CLV-cytokinin crosstalk is also an evolutionarily conserved feature of stem cell regulation. Genetic analyses reveal that CLV1 and RPK2 regulate SAM proliferation via separate pathways in moss. Surprisingly, cytokinin receptor mutants also form ectopic stem cells in the absence of cytokinin signaling. Through modeling, we identified regulatory network archtectures that recapitulated the stem cell phenotypes of *clv1* and *rpk2* mutants, cytokinin application, cytokinin receptor mutations, and higher-order combinations of these perturbations. These models predict that CLV1 and RPK2 act through separate pathways wherein CLV1 represses cytokinin-mediated stem cell initiation and RPK2 inhibits this process via a separate, cytokinin-independent pathway. Our analysis suggests that crosstalk between CLV1 and cytokinin signaling is an evolutionarily conserved feature of SAM homeostasis that preceded the role of WUS in stem cell organization.

## Introduction

Plant shoot morphology is generated by pluripotent stem cells called Shoot Apical Meristems (SAMs) at growing shoot tips. SAMs perform two essential functions during morphogenesis: to pattern lateral organ initiation and to maintain the stem cell population by replacing cells differentiated during organogenesis. Decades of study in the model flowering plant *Arabidopsis thaliana* have revealed a key role for a negative feedback loop in SAM homeostasis^1^. Stem cells at the apex of the meristem secrete a slew of CLAVATA3/EMBRYO SURROUNDING REGION (CLE) peptides including CLV3^2,3^. CLV3 acts through a suite of leucine-rich repeat receptor-like kinases (LRR-RLKs) including CLAVATA1 (CLV1) and RECEPTOR-LIKE PROTEIN KINASE2 (RPK2) to repress the expression of the stem cell organizing gene *WUSCHEL* (*WUS*), thereby limiting the size of the stem cell pool^4–8^. WUS promotes expression of *CLV3*, completing the negative feedback loop^9,10^. In Arabidopsis, *wus* mutants fail to maintain a SAM^11,12^. In contrast, *clv3, clv1*, or *rpk2* loss-of-function mutants fail to downregulate *WUS* expression and produce too many stem cells, leading to an enlarged and fasciated SAM, increased organ numbers, and stem swelling^1,4^.

Unlike the multicellular SAM of flowering plants, the moss *Physcomitrium patens* (henceforth Physcomitrium, previously known as *Physcomitrella patens*) SAM comprises a single tetrahedral stem cell^13^. Despite this anatomical difference, the moss apical cell accomplishes the same two essential SAM functions: lateral organ patterning and self-maintenance. In so doing, the moss SAM divides asymmetrically to form a leaf progenitor cell and a new apical stem cell. The apical division plane rotates to produce leaves (phyllids) in a spiral pattern around the haploid shoot (gametophore). The transcriptome of the moss SAM shares a high degree of overlap with flowering plant SAMs, suggesting possible deep homology underlying SAM function despite the fact that the moss haploid SAM and the diploid SAM of flowering plants reside in non-homologous shoots^14^.

In contrast to flowering plants, Physcomitrium encodes only three orthologs of stem cell-regulating LRR-RLKs. This relative simplicity makes moss an appealing model for understanding the role of these receptors in stem cell specification^15^. We previously characterized the function of *CLV1* and *RPK2* orthologs in Physcomitrium and found that these genes performed similar developmental functions in moss and moss^16^, including regulation of SAM homeostasis. Moss *clv1* and *rpk2* mutants produce ectopic stem cells, suggesting that the canonical roles of CLV1 and RPK2 to reduce stem cell number are conserved. Interestingly, whereas the Arabidopsis *clv1* and *rpk2* stem cell phenotypes are attributed to overaccumulation of WUS, *Physcomitrium patens* lacks this *WUS* function^17^. The Physcomitrium genome encodes three *WUSCHEL-RELATED HOMEOBOX* (*WOX*) genes, all of which lack domains critical for the stem cell modulatory function of WUS^17–19^. Moss *wox* mutants exhibit defective tip-growth in regenerating protonemal filaments, however, shoot development is normal^20^. Together, these data suggest that moss *WOX* genes do not function during SAM homeostasis^21^. This raises the question: in the absence of WOX-mediated stem cell maintenance, how do CLV1 and RPK2 inhibit stem cell specification in moss?

The hormone cytokinin promotes SAM formation in both Arabidopsis and Physcomitrium^30^. Several lines of evidence suggest an antagonistic relationship between CLV1 and RPK2, and cytokinin^16,22^. In moss shoots, cytokinin promotes the formation of new SAMs^23^, while loss of CLV1 or RPK2 signaling causes ectopic stem cell formation^16^. Intriguingly, in Arabidopsis, cytokinin promotes SAM formation via induction of *WUS* expression^24,25^. This again begs the question of how cytokinin promotes SAM formation in moss, despite the absence of WOX function during stem cell organization. In this study, we ask if there is crosstalk between CLV1, RPK2, and cytokinin signaling during stem cell specification in Physcomitrium.

Mathematical models representing competing hypotheses were fit to the empirical data on SAM homeostasis in wild type versus *clv1* and *rpk2* mutant shoots, with and without cytokinin treatment, and when cytokinin signaling is blocked. Maximum support was found for a model where CLV1 signaling is upstream of cytokinin-mediated stem cell induction, and RPK2 acts via a separate pathway. Overall, our data support a network in which CLV1 and cytokinin signaling converge in the regulation of moss SAM maintenance. This network is distinct from the canonical angiosperm network in that it lacks a role for a *WOX* gene as the hub linking cytokinin CLV1 pathways. Thus our work suggests crosstalk between CLV1-like receptors and cytokinin signaling is an evolutionarily-conserved feature of SAM homeostasis that preceded the role of WUS in stem cell organization.

## RESULTS

### CLV1 and RPK2 function through distinct pathways

Our previous research revealed a conserved role for the moss LRR-RLKs CLV1A, CLV1B, and RPK2 in inhibiting stem cell identity in moss shoots^16^. Loss of function mutations in the two *CLV1* paralogs or of *RPK2* resulted in shoots with ectopic apical cells along the lengths of swollen stems^16^. In order to determine whether the CLV1s and RPK2 function in the same pathway to regulate stem cell formation in moss, we used CRISPR-Cas9 mediated mutagenesis to target *RPK2* in a *clv1a clv1b* double mutant (hereafter designated as *clv1*) background and generated three independent, triple mutant lines with similar phenotypes (Supplemental Figure 1). We examined wild type, *clv1* double mutants, *rpk2* single mutants, and *clv1 rpk2* triple mutant shoots across three developmental timepoints ranging from less than one week old to approximately four weeks old (Figure 1).

**Figure 1:**
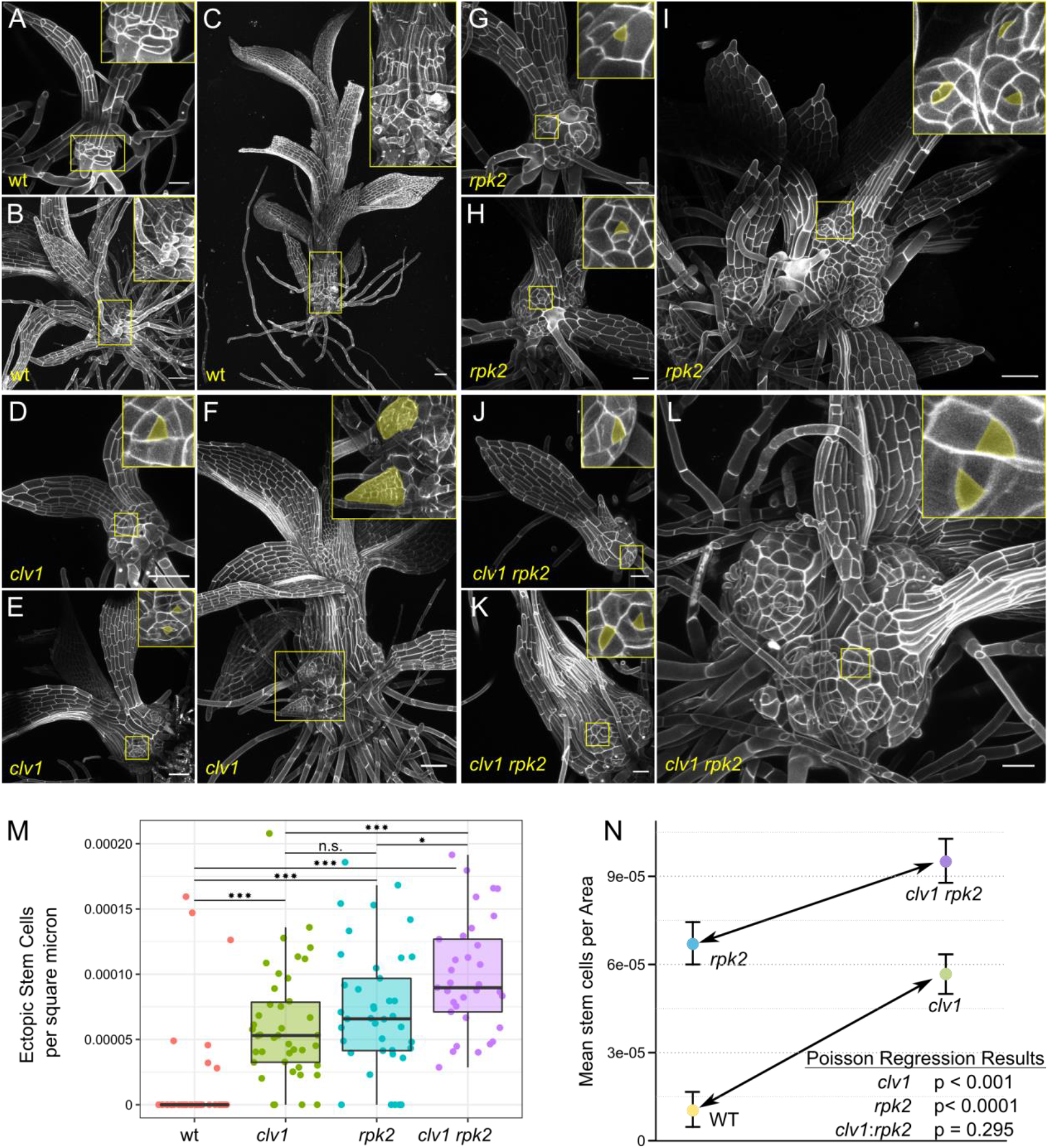
*clv1* and *rpk2 mutant* phenotypes are additive. Shoot development at approximately one week (A, D, G, J), two weeks (B, E, H, K) and three to four weeks (C, F, I, L) in wild type (A-C), *clv1* double mutants (D-F), *rpk2* (G-I), and *clv1 rpk2* triple mutants (J-L). Insets show cells at the base, which include ectopic stem cells and outgrowths initiating from these stem cells (pseudo colored yellow) in the mutants. (M) Ectopic stem cells and branches per square micron of visible stem tissue from 30 - 40 shoots of each genotype. One star denotes p < 0.05; three stars p < 0.001; two-sided t-test with Bonferroni correction for multiple testing. St (N) An interaction plot showing the effects of *clv1* (left to right, arrow) and *rpk2* (top to bottom) mutations on ectopic stem cell per square micron. The similar slopes of the two lines demonstrate that the effect of the two *clv1* mutations is the same in the wild type and *rpk2* genetic backgrounds. Results for a Poisson regression testing for significant effects from *clv1* double mutants, *rpk2* mutants, and an interaction between the two are given. Scalebar 50 µm.

Wild-type shoots were mostly covered with leaves, leaving little exposed stem tissue (Figure 1 A, B, C). In contrast, the stems of *clv1* double mutants, *rpk2* single mutants, and triple mutants were swollen from the earliest stages of development (Figure 1D, G, J). Ectopic apical cells were present on the rounded stems of these mutants at all stages of development. Apical cells were identified by the characteristic triangular shape of their apical surface, the top of the tetrahedron. At later stages, *clv1* double, *rpk2*, and *clv1 rpk2* triple mutant shoots produced ectopic leaves, indicating that the ectopic apical cells observed at earlier stages were indeed functional SAMs (Figure 1F).

All mutant shoots stopped elongating earlier than the wild type, often terminating in a swollen apex with abundant apical cells (Figure 2). After terminating longitudinal growth, *clv1* double, *rpk2*, and *clv1 rpk2* triple mutant shoots continued to swell and initiate new growth axes from ectopic stem cells along the length of the stem (Figure 1 F, I, L, Figure 2 B). The swelling of shoot tips in a mass of stem cells in *clv1* and *rpk2* mutant stems was reminiscent of stem cell over-proliferation and SAM disorganization seen in Arabidopsis *clv* and *rpk2* mutants^4,26^, again highlighting the conserved function of this pathway in SAM homeostasis^16^.

**Figure 2:**
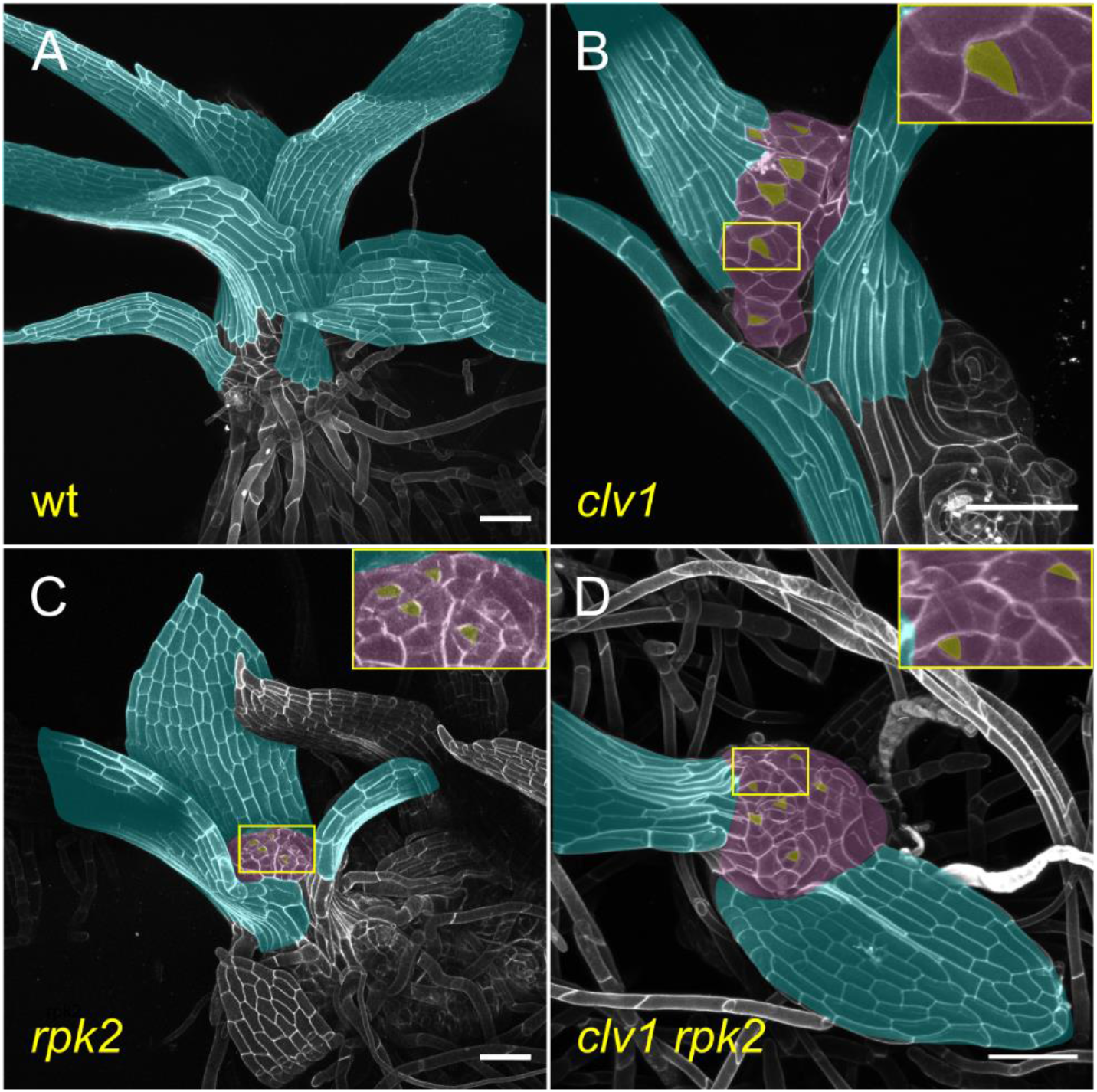
*clv1* and *rpk2* shoots terminate in a mass of stem cells. Approximately three-week-old wild type (A), *clv1* double (B), *rpk2* (C), and *clv1 rpk2* triple mutant (D) shoots. Bare shoot apices (pseudo colored purple) replete with stem cells (pseudo colored yellow) are visible in *clv1* double, *rpk2* and *clv1 rpk2* mutants, while the single apical cell at the apex of wild type shoots is well-covered by leaves (pseudo colored blue). stem cell Scalebars: 100 nm.

However, it was not clear how CLV1 and RPK2 signaling interact to inhibit ectopic stem cell formation. To determine whether CLV1 and RPK2 function in the same linear pathway, we quantified the number of visible ectopic stem cells by confocal microscopy on 30 - 40 shoots of each genotype. To control for variation in age and any increase in stem cell count due to increased mutant stem size, ectopic stem cell number was normalized to the size of the stem (Figure 1 M, non-normalized data Supplemental Figure 2). Ectopic stem cells were observed on almost all *clv1* and *rpk2* mutants, in contrast with wild type.

If *CLV1* and *RPK2* act in a linear genetic pathway, epistasis is predicted in *clv1 rpk2* triple mutants. In contrast, if CLV1 and RPK2 have redundant functions but act in distinct complexes, a synergistic increase in stem cell initiation is predicted in triple mutants. Finally, if CLV1 and RPK2 act non-redundantly, i.e. in distinct signaling pathways, additive effects on stem cell initiation phenotypes are expected. The number of ectopic stem cells per area was significantly higher in *clv1 rpk2* triple mutants than either *clv1* doubles (p < 0.001) or *rpk2* single mutants (p = 0.01) (Figure 1 M). An interaction plot revealed that the effects of mutating *clv1a clv1b* in wild type or *rpk2* mutant backgrounds (the slope of the arrow-headed line) is similar in both genotypes; thus, the effects of the *clv1* and *rpk2* mutations appear additive (Figure 1 N). A Poisson regression, which is suited for low count data and included stem area as an offset to control for stem size, was used to assess the statistical significance (p-value) and impact (coefficient) of mutating *CLV1* and *RPK2* and the interaction between these two (Figure 1 N). The regression revealed significant effects for loss of CLV1 and RPK2 function (p < 0.0001 for each and coefficients 0.5054 and 0.6358 for *clv1* and *rpk2*, respectively), but no significant interaction between them (p = 0.29, coefficient -0.1774). In summary, the genetic data suggest that CLV1 and RPK2 do not function in the same linear pathway during regulation of stem cell abundance in the moss shoot.

### CLV1 and RPK2 signaling interact with Cytokinin signaling

Exogenous cytokinin application induces swelling and the stem cell formation along wild-type moss shoots^23^, similar to phenotypes observed in *clv1* and *rpk2* mutants (Figure 3A-C and Figure 1). We hypothesized two possible pathways to explain the convergence of these phenotypes. First, CLV1 and/or RPK2 could function by inhibiting cytokinin-mediated stem cell specification, i.e. CLV1/RPK2 function is upstream of cytokinin response. Alternatively, cytokinin signaling might be upstream and inhibit CLV1/RPK2 function, such that cytokinin de-represses stem cell formation. To better understand how CLV1 or RPK2 signaling interact with cytokinin signaling, we characterized the response of *clv1* double, *rpk2*, and *clv1 rpk2* triple mutants to exogenous treatment (10 nM and 100 nM) of the synthetic cytokinin 6-benzylamino purine (BAP). If CLV1 or RPK2 function upstream or independently of cytokinin signaling, BAP treatment is predicted to induce stem cell formation in *clv1* and *rpk2* mutants. On the other hand, if cytokinin is an upstream inhibitor of CLV1 and/or RPK2 function, BAP treatment is expected to have no effect on the *clv1* or *rpk2* mutant phenotypes.

**Figure 3:**
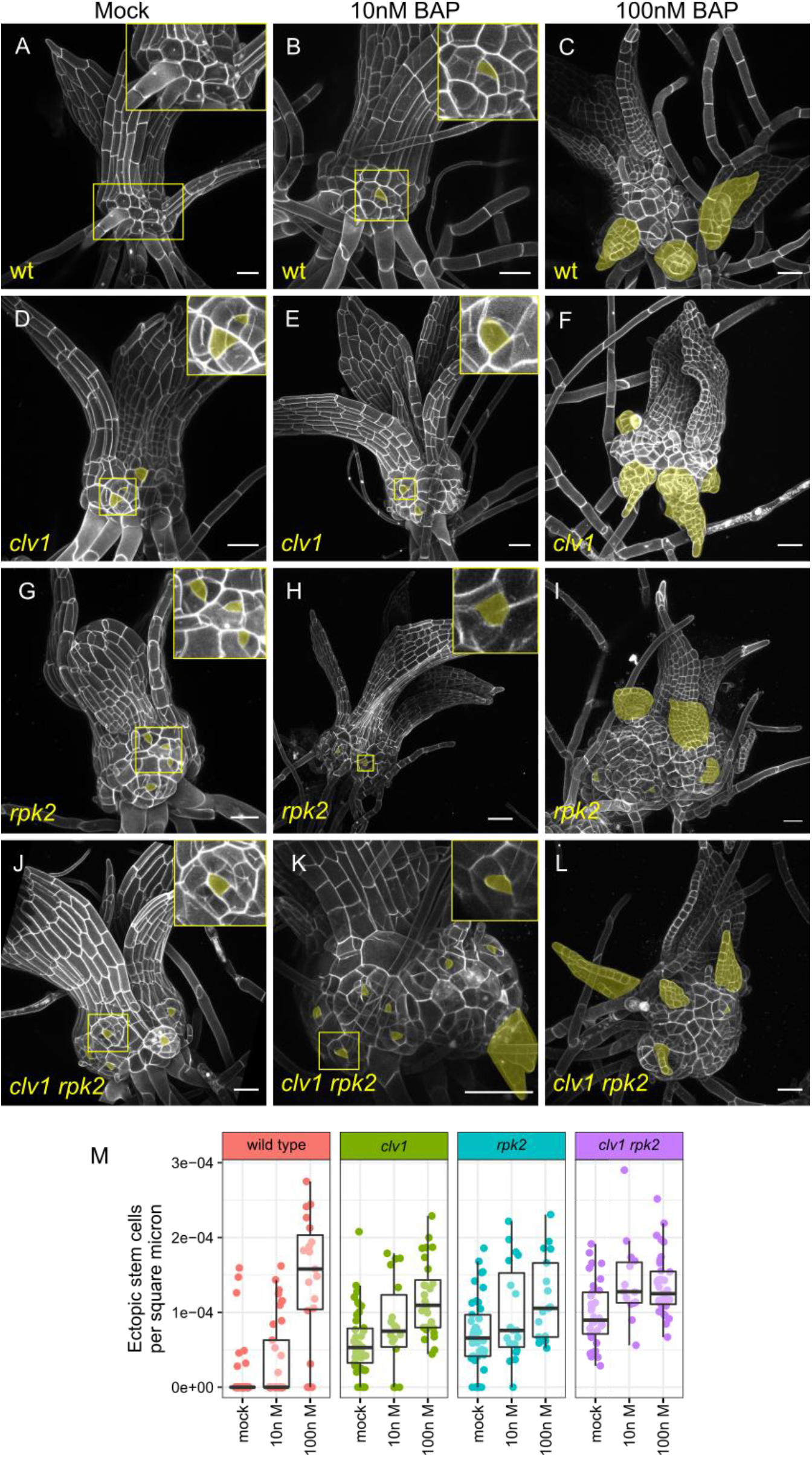
Cytokinin induces phenotypes similar to *clv1*/*rpk2* and increases apical cell formation in *clv1/rpk2* mutants. Two-to three-week-old shoots grown on mock media (0 nM A, D, G, J) or media supplemented with low (10 nM, B, E, H, K) and high (100 nM, C, F, I, L) BAP. (M) Quantification of ectopic stem cells per area of exposed stem. Note that (M) includes data from Figure 1M for clarity. Ectopic stem cells and ectopic leaves (indicative of underlying stem cells) are pseudo colored yellow. Scalebars: 50 µm.

We quantified the number of ectopic stem cells in each genotype grown on 10 nM and 100 nM cytokinin, normalized to stem area as described above. Wild-type shoots grown on 10 nM BAP displayed slight swelling and developed ectopic tetrahedral apical cells reminiscent of those seen in *clv1* and *rpk2* mutants (Figure 3A-B). A higher concentration of cytokinin rendered swollen stems and numerous outgrowths derived from ectopic stem cells in wild-type plants (Figure 3C, M). In comparison, 10 nM BAP had mild effects on stem swelling and ectopic apical cell formation in *clv1* and *rpk2* mutant shoots (Figure 3 E, H, M), although treatment with 100 nM BAP induced numerous ectopic apical cells and a high degree of stem swelling (Figure 3F, I, M). Interestingly, growing *clv1 rpk2* triple mutants on 10 nM or 100 nM BAP resulted in a similar amount of stem cells per area (Figure 3 K, L, M, non-normalized data Supplemental Figure 3), suggesting ectopic stem cell formation was already saturated. Overall, stem cell formation increased in *clv1* double, *rpk2*, and *clv1 rpk2* triple mutants in response to cytokinin. These data support a role CLV1 and RPK2 upstream or independent of cytokinin response.

Interestingly, while cytokinin induced stem cell formation in all genotypes, this effect appeared weaker in *clv1* and *rpk2* mutants than in wild type (Figure3 M). To statistically assess how CLV1, RPK2, and cytokinin interact to control stem cell specification, we analyzed our full dataset, comprising each genotype with and without BAP treatment, using a Poisson regression. Notably, there remained a significant increase in stem cells caused by loss of *CLV1* or *RPK2* function, as well as cytokinin treatment (p-value and Poisson coefficient for *clv1*: p = 0.0001, coefficient = 0.64; *rpk2*: p < 0.0001, coefficient = 0.96; cytokinin treatment: p < 0.0001, coefficient = 0.0133, see Materials and Methods for details). In agreement with our conclusion from Figure 1, cytokinin treatment also revealed no significant statistical interactions between the effects of *clv1* and *rpk2* mutations (p = 0.240), indicating that these mutant phenotypes are indeed additive. Interestingly, each mutant showed a slight but statistically significant reduced induction of stem cells in response to exogenous cytokinin compared to wild type (Figure 2I; *clv1*:cytokinin p = 0.033, coefficient = -0.003; *rpk2*:cytokinin p < .0001; coefficient = -0.009;).

In summary, the phenotypes of *clv1* and *rpk2* were additive across a range of exogenous cytokinin concentrations, suggesting that CLV1 and RPK2 act via distinct pathways regulating stem cell specification. Cytokinin increases stem cell initiation in *clv1* and *rpk2* mutants, supporting a role for CLV1 and RPK2 upstream or independent of cytokinin signaling. However, loss of *CLV1* or *RPK2* function also slightly diminished the effect of exogenous cytokinin on stem cell production. These data suggest that cytokinin response could already be high in *clv1* and *rpk2* mutants such that exogenous cytokinin treatments represented a smaller relative increase in cytokinin signaling in the mutants than in wild type. Such a scenario would occur if CLV1 and/or RPK2 functioned by inhibiting cytokinin-mediated stem cell specification.

### Mathematical modelling supports action by CLV1 and RPK2 through distinct pathways

To test possible regulatory network topologies that integrate CLV1, RPK2, and cytokinin signaling to control stem cell identity in the moss shoot, we evaluated a range of hypothetical gene regulatory network models (e.g. Figure 4 A). Given that the data supported CLV1 and RPK2 signaling through separate pathways, we coded two variables, ‘x’ and ‘y’, to represent two hypothetical pathways each capable of promoting stem cell formation. It is important to note that x and y do not represent any specific genes, but rather are simplified representations of unknown but postulated signaling outputs. As cytokinin induces stem cell formation, we specified that pathway x denotes cytokinin response, while pathway y represents cytokinin-independent stem cell specification pathways.

**Figure 4:**
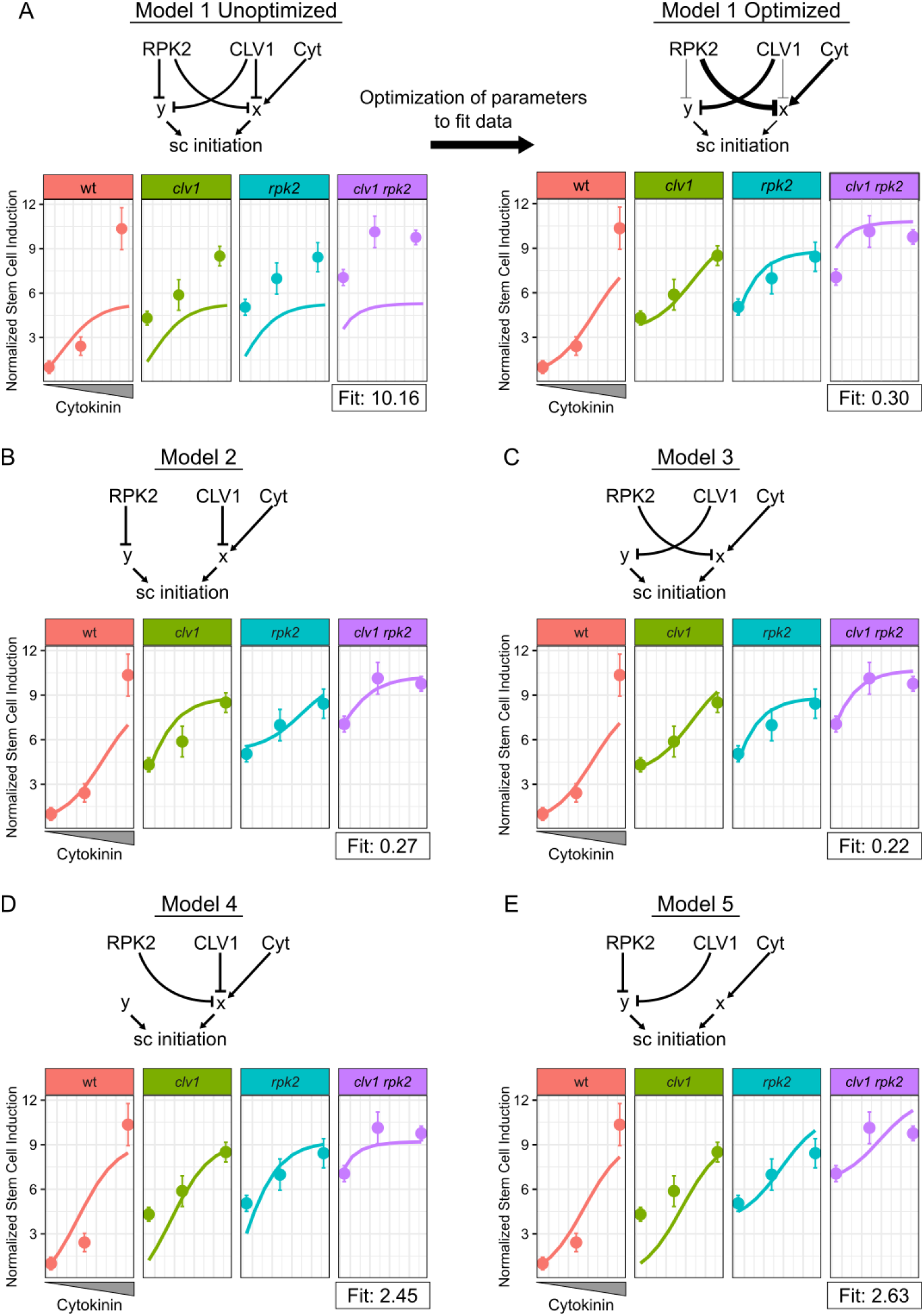
Dynamical model simulations of stem cell production by *clv1* and *rpk2* mutants over a range of cytokinin concentrations. Basic network depictions and simulation results of mathematical models formalizing five different hypotheses: Model 1 CLV1 and RPK2 are partially redundant (A); Model 2 CLV1 and RPK2 are independent with CLV1 upstream of cytokinin response (B); Model 3 CLV1 and RPK2 are independent with RPK2 upstream of cytokinin response (C); Model 4 CLV1 and RPK2 are redundant and upstream of cytokinin response (D); Model 5 CLV1 and RPK2 are redundant and upstream of a cytokinin-independent pathway (E). On plots, Solid lines represent simulated data over a range of cytokinin concentrations. Dots with error bars represent the mean empirical stem cells per square micron data and standard errors. From left to right, dots represent values from moss grown on mock, 10 nM BAP, and 100 nM BAP. The model on the right side of panel A has had the thickness of interactions changed proportional to the corresponding optimized parameters. Note the empirical data points are based on the data in Figure 3M.

In the first model, RPK2 and CLV1 are capable of inhibiting stem cell initiation through both cytokinin-dependent and independent pathways, i.e. through both x and y (Figure 4 A). This model has the greatest flexibility and represents a scenario where CLV1 and RPK2 have overlapping but non-identical contributions to both the x and y pathways. The second and third models simulate the cases where CLV1 and RPK2 act completely independently (Figure 4 B and C). In model two CLV1 inhibits cytokinin-mediated stem cell induction (x) and RPK2 inhibits cytokinin-independent stem cell induction (y), while the roles for CLV1 and RPK2 are reversed in model three. As an additional test of whether *CLV1* and *RPK2* might be redundant, we included models four and five that wherein each inhibits x (Figure 4 D) or y (Figure 4 E) exclusively, although with potentially different strengths. Thus, these five models represent five competing hypotheses where CLV1 and RPK2 are partially redundant, completely independent, or completely redundant, and where each acts upstream of a cytokinin-dependent or a cytokinin-independent pathway promoting stem cell specification.

We next compared the extent to which each model could recapitulate the patterns of ectopic stem cell production seen in the empirical data. The behavior and output of a model depends on the parameters selected. Thus, we sought to optimize the parameters in each network to reach the best fit possible to the empirical data. To find the optimal parameters, we coded a random optimizer function (see Methods). Given a network and a set of starting parameters, we used the model to simulate each relevant mutant genotype (wild type, *clv1*double, *rpk2*, and *clv1rpk2* triple mutants) at each level of cytokinin treatment (0 nM BAP, 10 nM BAP, 100 nM BAP). For each model we thus simulated twelve scenarios that we compared with the corresponding mean values from the empirical data. We generated a single fit score that was proportional to the difference between the model and the empirical data (smaller scores indicate a closer fit). Larger differences between simulated and empirical data points were penalized more heavily than smaller ones (See Model Methods). Once a score was generated, each parameter was randomly mutated (adjusted up or down) and a new fit score was calculated and compared to the previous one. If the new fit score was better (lower), the new parameters for that run were adopted as the starting point for the next round of optimization. If the fit score proved worse, the previous parameters were kept as the starting point for the next round of optimization. We ran the optimizer for 300 iterations, allowing the fit score to plateau at a minimum value for each model (see Model Methods).

Using this iterative optimizer, we tested our five competing models of how CLV1, RPK2, and cytokinin regulate stem cell identity. For model one, the optimizer selected parameters that separated the roles for CLV1 and RPK2, i.e., the optimizer minimized their redundant activity and emphasized their independent activity (Figure 4 A). Specifically, the optimizer strengthened the regulatory connection between RPK2 and cytokinin-dependent signaling while weakening the interaction between RPK2 and cytokinin-independent induction, and the optimizer did the opposite for CLV1 (Figure 4 A). Thus, the optimized Model 1 was similar to Model 3 (Figure 4 C). Optimized Model 1 reasonably replicated the empirical data with a fit score of 0.3, but notably overestimated the number of ectopic stem cells in the triple mutant grown in the absence of exogenous cytokinin (Figure 3 A). Models 2 and 3 produced nearly equivalent fits to Model 1 after optimization with fit scores of 0.27 and 0.23, respectively (Figure 5 B, C). Notably, Model 2 and Model 3 successfully simulated the level of stem cell initiation in triple mutants without exogenous cytokinin. These two models differed from one another predominately in the cytokinin response curves of *clv1* and *rpk2* mutants (Figure 3 B and C), but neither simulation was far from the empirical data. Finally, Model 4 (fit score = 2.45) and 5 (fit score = 2.63) produced very poor fits to the data (Figure 2 D and E), highlighting the unlikelihood of redundancy between CLV1 and RPK2. Altogether, the five models support CLV1 and RPK2 acting through separate pathways. However, the models do not confidently distinguish whether

**Figure 5:**
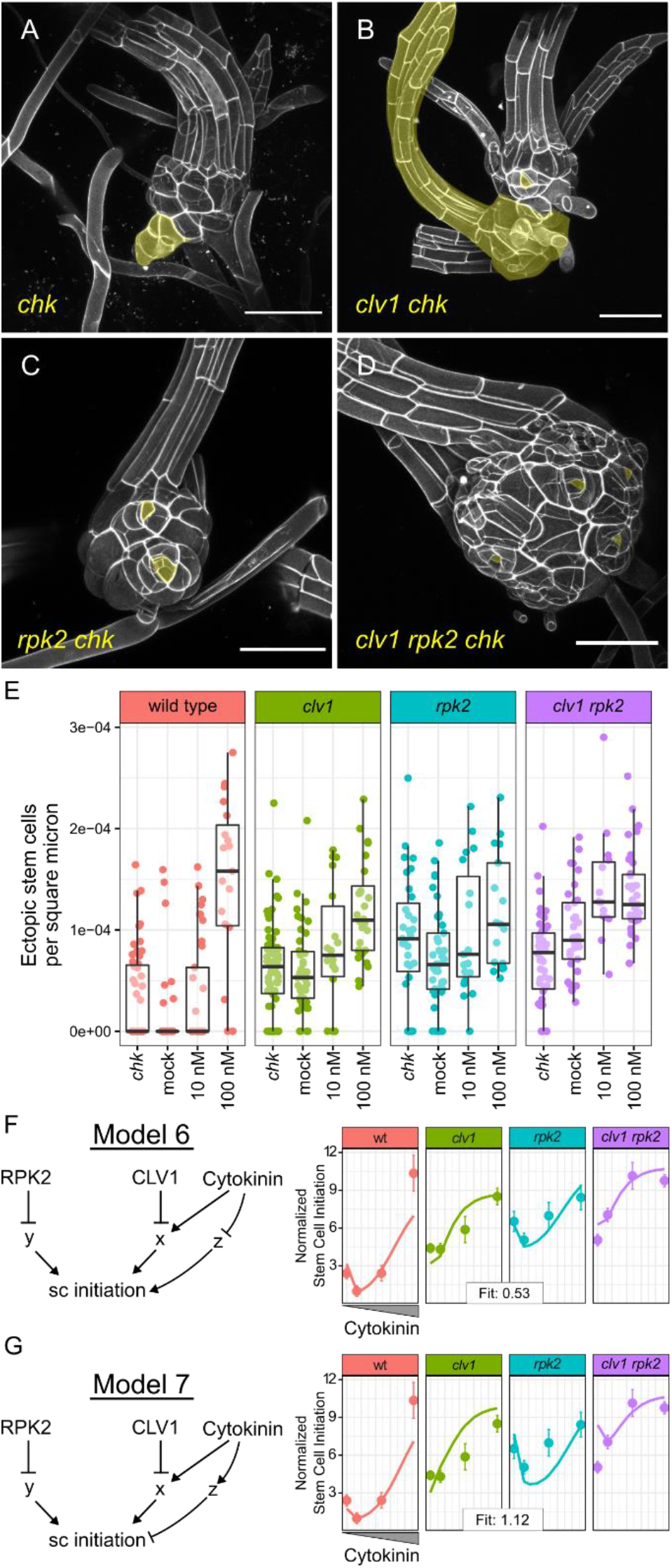
*chk* mutants produce ectopic stem cells and have complex interactions with *clv1* and *rpk2* mutants. Shoots from five week old colonies of *chk* (A), *clv1a clv1b chk* quintuple mutants (B), *rpk2 chk* quadruple mutants (C), and *clv1 rpk2 chk* sextuple mutants (D). Ectopic stem cells or derived outgrowths highlighted in yellow. Quantification of ectopic stem cells per square micron of visible stem (E), including data from Figure 2 I. Model six posits that cytokinin inhibits an inducer of stem cell identity (F). In model seven, cytokinin promotes an inhibitor of stem cell specification (G). In (F) and (G), solid lines represent simulated data while dots represent mean stem cells per area from the empirical data. Error bars show the standard error. The x axis shows a log transformation of the cytokinin value input to the model.

CLV1 or RPK2 functions upstream of the cytokinin response as either scenario could reproduce the empirical data.

Given that models 2 and 3 best reproduce the empirical data”, we used them to predict how stem cell initiation would be affected by the absence of cytokinin signaling (Supplemental Figure 4). Each model predicted that if devoid of cytokinin signaling, every simulated genotype would produce even fewer apical cells. This is consistent with data showing reduced branch stem cell initiation after increasing cytokinin degradation^23^. Informatively, the model predicted that at zero cytokinin perception, ectopic stem cell formation resulting from loss of whichever receptor (RPK2 or CLV1) that functioned upstream of cytokinin dependent stem cell initiation would be suppressed (Supplemental Figure 4).

### Loss of cytokinin signaling reveals a complex network controlling stem cell specification

Three *CYTOKININ HISTIDINE KINASE* (*CHK*) genes in Physcomitrium encode the known cytokinin receptors, and *chk1 chk2 chk3* triple mutants (hereafter designated as *chk*) lack the ability to perceive cytokinin^27,28^. Using CRISPR-Cas9 to mutate *CLV1a, CLV1b*, and *RPK2* in the *chk* background, we generated and confirmed four independent lines of *clv1 chk* quintuple mutants, two independent *rpk2 chk* quadruple mutants, and two *clv1 rpk2 chk* sextuple mutants (Supplemental Figure 5). Our models predicted that if ectopic stem cell formation in *clv1* mutants resulted from increased cytokinin-mediated stem cell initiation, ectopic stem cell formation would be suppressed in the higher order *clv1 chk* mutants. Alternatively, if RPK2 signaling were upstream of the cytokinin response (x), the *rpk2* ectopic stem cell phenotype would be suppressed in *rpk2 chk* mutants. To test our predictions of how loss of cytokinin signaling impact stem cell specification, we examined shoots from multiple independent lines of *chk* and higher order mutants at cellular resolution and quantified stem cell abundance (Figure 5, A-E).

Cytokinin induces the formation of shoots^23^ and promotes cell proliferation in leaves (Figure 3A-C)^29^. Consistent with these functions, the *chk* triple mutants develop shoots two weeks later than wild-type moss^28^. Once formed, *chk* shoots produce long, slender leaves with highly elongated cells and fewer cell files (Supplemental Figure 6). Surprisingly, confocal imaging of *chk* mutant shoots revealed the formation of ectopic growth axes, indicating increased stem cell production. Higher order *clv1 chk* quintuple, *rpk2 chk* quadruple, and *clv1 rpk2 chk* sextuple mutants produced swollen stems with variable degrees of ectopic apical cells (Figure 5 A-E; Supplemental Figure 6, non-normalized data Supplemental Figure 7). We statistically analyzed the full dataset including stem cell quantification from all mutants and genotypes generated thus far using a Poisson Regression. We found that loss of CHK function was significantly associated with increased stem cell abundance (p = 0.013). Although neither loss of CLV1 or RPK2 displayed statistically significant interactions with CHK loss of function, a negative interaction between *clv1* and *chk* mutations tended towards significance (*clv1:chk* p = 0.07; *rpk2:chk* p = 0.11), weakly supporting a role for *CLV1* upstream of cytokinin. Overall, loss of cytokinin signaling causes ectopic stem cell formation and does not suppress stem cell initiation, contrary to our predictions. Thus, the *chk* mutant phenotypes suggest that important features were missing from our dynamical models of stem cell homeostasis in moss.

### Incoherent feedforward control of stem cell induction can explain *chk* phenotypes

We hypothesized that the ectopic stem cell phenotype of *chk* triple mutants could be explained by an incoherent feedforward control: in addition to its primary function promoting stem cell formation, cytokinin either (1) represses a pathway that induces stem cell specification, or (2) promotes a pathway that represses stem cell specification. To test the plausibility of these hypotheses, we formalized each as a dynamical model (Model 6 and 7, respectively), each with versions based on Models 2 and 3. For each model, we introduced a factor called z, which is independent of CLV1 and RPK2 signaling. In Model 6, z induced stem cell initiation and was inhibited by cytokinin (Figure 5 F), while in Model 7 z inhibited stem cell initiation and was itself induced by cytokinin (Figure 5 G). Each model represents an alternative form of the hypothesis that cytokinin promotes stem cell induction but also provides additional input to temper the sensitivity to stem cell specification cues. We predicted that in the complete absence of cytokinin signaling, the loss of this buffering capacity would render stem cell specification hypersensitive to inputs from other pathways, such as those represented by y in the model.

Optimizing each model to the full dataset provides a test of how well each model can capture key trends in the empirical data. Because we previously found that either CLV1 *or* RPK2 acted upstream of cytokinin signaling (x), we simulated both scenarios for each model (Figure 5 F, G, Supplemental Figure 8). Model 6 successfully reproduced the trends in the empirical data, and fit better if CLV1 were upstream of cytokinin response (fit score = 0.53, Figure 5F) rather than RPK2 (fit score = 0.74, Supplemental Figure 8). Model 7 - in which cytokinin induced an independent stem cell inhibitory pathway – fit the data poorly in all cases as it overestimated the number of ectopic stem cells forming in higher order *clv1, rpk2* and *chk* mutants (Model 7 fit scores: 1.12 with and 1.36 with CLV1 and RPK2 upstream of cytokinin, respectively) (Figure 5 G, Supplemental Figure 8). In comparison, when model 2 and model 3 were fit to this dataset they produced fit scores of 1.85 and 1.59, respectively (Supplemental Figure 9), demonstrating the impact of including ‘z’ in the model. Overall, a role for cytokinin in buffering stem cell initiation can explain the ectopic stem cell formation in *chk* mutants and higher order *clv1 chk* and *rpk2 chk* mutants.

## Discussion

Our work demonstrated similar functions for CLV1 and RPK2 as regulators of stem cell abundance in the moss Physcomitrium as have been previously reported in Arabidopsis. We used a combination of higher-order genetics, hormone treatment, and mathematical modeling to demonstrate that, as in flowering plants, stem-cell identity in moss shoots is regulated by interaction between CLV and cytokinin signaling.

### CLV1 and RPK2 signal through distinct pathways

In Arabidopsis, it is unclear whether CLV1 and its paralogs function in the same pathway as RPK2^3031,32^. The suite of CLV1 and RPK2-like receptors in moss is much reduced compared to Arabidopsis, making it a powerful system for studying their genetic interactions^16^. *clv1* and *rpk2* have distinct filament phenotypes in moss, where *rpk2* colonies spread faster than wild type or *clv1* colonies^16,33^. This phenotypic distinction could arise from differences in expression patterns, rather than distinct molecular functions. However, *CLV1* and *RPK2* are co-expressed in shoots, where their loss of function render similar mutant phenotypes^16^. Here we show that CLV1 and RPK2 act additively in shoots, in distinct pathways, to regulate stem-cell specification.

### Cytokinin CLV crosstalk in moss and Arabidopsis have similar networks, except for WUS

In Arabidopsis, CLV1, RPK2, and cytokinin regulate stem-cell identity as part of a network centered around the master regulator gene *WUSCHEL*. In moss, *WOX* genes do not regulate SAM homeostasis^26^. Both exogenous cytokinin and decreased CLV1/RPK2 function cause similar shoot phenotypes, including stem swelling and ectopic stem-cell formation. This overlap in phenotypes led us to ask whether CLV1/RPK2 and cytokinin interact to regulate stem-cell identity in moss. We tested the response of *clv1, rpk2*, and *clv1 rpk2* triple mutants to cytokinin and loss of cytokinin signaling (*chk)* and used mathematical modeling to determine which of seven different hypothetical networks describing CLV1, RPK2, and cytokinin function could best recapitulate empirical stem-cell specification data. Our data and modeling suggest that either CLV1 or RPK2 acts upstream of cytokinin-mediated stem-cell induction, with greater support for CLV1 performing this role. Our models represent overarching signaling networks, where x and y represent whole signaling cascades, not single genes. Thus, a number of specific molecular networks could fall within the frameworks of the model. Interestingly, this cytokinin-dependent pathway (x in the model, Figure 4, Figure 5) occupies the same position as *WUS* in models describing stem-cell specification in the Arabidopsis SAM^1,34^. This suggests that in the absence of *WUS*, stem-cell abundance in moss is regulated by similar mechanism as described in Arabidopsis, although *WUS* function is replaced by some unknown factor(s).

### Cytokinin signaling both induces and inhibits SAM formation

Our analysis of *chk* triple mutants and higher order *clv chk, rpk2 chk*, and *clv rpk2 chk* mutants revealed several unexpected phenotypes. First, we observed that *chk* mutants make ectopic stem cells, contrasting with cytokinin’s role promoting stem cell formation. We proposed several models to explain the counterintuitive *chk* and higher-order mutant phenotypes. An incoherent feedforward model where cytokinin signaling buffers stem cell initiation through a CLV and RPK2-independent pathway successfully replicated the trends seen in the empirical data, over a wide range of mutant genotypes and hormone treatments. However, because little is known about genes promoting SAM formation in moss, it is difficult to speculate as to the nature of ‘z’ in the model. It is possible that auxin signaling, which is upstream of stem cell formation on filaments and shoots, is de-regulated in *chk* mutants^23,35^. However, cytokinin application does not modulate stem cell-inducing factors downstream of auxin^35,36^. Thus, elucidating the identity of ‘z’ requires better understanding of cytokinin-responsive genes in moss shoots.

### Common molecular mechanisms can underlie disparate developmental functions across plant evolution

Similarities and differences in stem-cell regulation between moss and Arabidopsis raise important questions about the evolution of this signaling network. Current phylogenies support a monophyletic clade containing mosses and liverworts, such that mosses and liverworts are equally related to flowering plants. However unlike in moss and Arabidopsis, CLV1 function in Marchantia enhances stem-cell specification^37^. Interestingly, CLV1-mediated control of the meristem in Marchantia is also independent of *WOX* genes, supporting the later recruitment of *WOX* genes to regulation of the SAM^37^. Since SAM specification occurs in a *WUS*-independent pathway in bryophytes, is there also a *WUS-*independent pathway in angiosperms? Overall, it is curious how the moss SAM transcriptome is highly enriched for orthologs of genes that function in angiosperm SAMs, but lacks a role for *WOX* genes, which establish the canonical connection between CLV and cytokinin signaling in flowering plants. These observations raise the question of whether similarities between moss and flowering plant stem-cell specification pathways are convergent or whether CLV-cytokinin interactions are ancient with *WOX* genes incorporated with the evolution of the multicellular SAM.

### Brief Materials and Methods

For detailed materials and methods, see Supplemental Methods. Gransden 04 *Physcomitrium patens* protonema was propagated on BCDAT (see Media) for routine culture, while for phenotyping, moss was grown on BCD. Moss transformation was performed using a modified protocol described previously (see supplemental methods for detailed protocol) ^16,38^. gRNAs were expressed using the moss U6 and U3 promoters; Cas9 expression was driven under the rice Actin promoter^39^. Imaging was conducted using a Zeiss 710 confocal microscope; moss was stained for 15 minutes with Propidium Iodide prior to imaging. A 514 nm laser was used for excitation, and 566 to 650 nM were collected for emission. Ectopic stem cell counts and stem areas were measured in FIJI^40^ using maximum intensity projections or full z-stacks. Statistical analyses were performed in R: Poisson models were coded as general linear models set to the ‘Poisson’ family and included the log of the area as an offset.

## Acknowledgements

We would like to thank the Cornell Statistical Consulting unit and Celine Cammarata for advice on statistical analyses and data processing. We also thank Jill Harrison, Zoe Nemec Venza for thoughtful and helpful discussions about this work. We thank Jill Harrison, Zoe Nemec Venza, Mingyuan Zhu, Kate Harline, and Shuyao Kong for comments on the manuscript. This research was supported by NSF IOS-1238142 to M.J.S. and NSF IOS-1553030 to A.H.K.R., the Schmittau-Novak Small Grant to J.C., the Cornell Provost Diversity Dissertation Completion Fellowship to J.C., funding from the Weill Institute for Cell and Molecular Biology to A.H.K.R., and the NSF funded Boyce Thompson Institute Cornell REU program support for C.M.F (REU #1850796).

