## supplemental methods for "Cytokinin-CLAVATA crosstalk is an ancient mechanism regulating shoot meristem homeostasis in land plants"

#### Moss Culture

The Gransden 04 strain of *Physcomitrium patens* (formerly *Physcomitrella patens*) was used for all experiments[1]. To propagate moss, protonemal tissue was blended in 5-7 ml sterile water using a dremel with a custom propeller blade attachment, and 1-2 ml of moss tissue was inoculated onto BCDAT (see media supplemental table) plates overlain with sterile cellophane under continuous light at 25 degrees Celsius. For phenotyping gametophores, small samples of freshly blended protonema were placed onto BCD agar media supplemented with the specified amount of filter sterilized BAP. These tufts of tissue grew for the specified time before gametophores were dissected for imaging.

#### Moss Transformation

A modified moss transformation protocol was used to permit transformation with smaller amounts of DNA and tissue[2,3]. Protonema grown for 5-7 days on BCDAT (2-4 plates) was placed into 15 ml 8% Mannitol solution in a petri dish. 5 ml of 2% Drisilase solution (see solutions table) was then added, and the dish was placed on a shaker set to 80 RPM for one hour. Next, the solution was gently pipetted through a 70 µm mesh into a 50 ml conical tube. Filtered protoplasts were spun at 250 g for five minutes with low brake. The liquid was then poured off and replaced with 20 ml 8% mannitol, and this wash step was repeated two more times. Upon being resuspended in mannitol solution a third time, the concentration of protoplasts was measured using a hemocytometer. Cells were then pelleted one final time and resuspended in a volume of 3M solution (solutions and media) to yield $2*10^6$ protoplasts/ml. 300 ul of the protoplast/3M solution was transferred to a 2 ml tube with a maximum of 10 ug DNA (with equimolar ratios of each plasmid, including each gRNA-encoding plasmid, the Cas9 expression plasmid, and selectable marker), and stirred gently. The DNA, 3M, and protoplast mixture was then transferred to a 15 ml conical tube with 300 ul PEG solution inside it; these were mixed gently by pipetting. The DNA and protoplast solution was heat shocked at 45 degrees Celsius for three minutes and then left to cool in a bath of room temperature water for 10 minutes. The solution was slowly diluted to a final volume of 5 ml with 8% mannitol over a period of half an hour, and then spun down to pellet the protoplasts. During the previous step, PRMT (solutions and media) was melted, supplemented with $CaCl_2$ to a final concentration of 10 mM and left to keep warm in a 45 degree water bath. PRMB (solutions and media) plates were overlain with sterile cellophane, pelleted protoplasts were resuspended in PRMT, and then 1 ml of the PRMT and protoplast mixture was plated onto each PRMB plate.

#### Moss Selection

Seven days after transformation, the cellophane with transformed protoplasts was transferred off the PRMB plates and onto BCDAT plates supplemented with 20 mg/L G418. After one week to ten days on selection, regenerating colonies were transferred to new BCDAT plates and left to grow until individual colonies were visible by eye (two-to-three weeks). Individual colonies were transferred to BCDAT plates without cellophane, and left to grow for three more weeks. After this time, these colonies were split and samples were taken for DNA extractions and genotyping by PCR and Sanger sequencing.

#### Plasmid Construction

For routine cloning of gRNAs into a transient expression vector suitable for use in *P. patens*, we used the pENT-U6pro::Bsa1:sgRNA and pENT-U3pro::Bsa1:sgRNA constructs generated previously[3]. To clone gRNA expression vectors, oligonucleotides with overhangs complementary to the BSAI cut sites (GGC and CAT for U3 and U6 promoters, and aaac to ligate on the 3' end with and were annealed. pENT-U3pro::Bsa1:sgRNA or pENT-U6pro::Bsa1:sgRNA were digested with BSAI, and linearized plasmids were purified by gel extraction. Annealed oligonucleotides were then ligated into the linearized gRNA vectors using T4 DNA ligase. Ligated DNA constructs were transformed into DH5-alpha competent cells grown in the presence of 50 µg/ml kanamycin. Clones were screened by colony PCR using the forward oriented gRNA oligonucleotide and pJCm208 (in the gRNA scaffold) as primers followed by Sanger sequencing using the pJCm208 primer. gRNA sequences were selected using the CRISPOR online service[4]. We minimized off target effects and where possible chose gRNAs that cleaved in the first exon. We tried four gRNAs for the *CLV1a* gene before finding one that cleaved effectively (not shown); otherwise the first gRNAs designed for each gene successfully induced mutations.

#### Genotyping

For low numbers of moss samples, tissue was ground by hand, while for large numbers of samples we purified DNA in 96 well plates. For small DNA preps and some 96 well-plate preps, tissue was ground in water, to which an equal volume of 2x Shorty Extraction buffer (0.4 M Tris, pH 9.0; 0.8 M LiCl; 50 mM EDTA; 2% SDS) was added. The debris were pelleted and the supernatant removed to a new tube to which an equal volume of isopropanol was added. The solution was then mixed by inversion and placed on ice or at -20 degrees Celsius for 15-30 minutes and then spun at maximum speed to pellet the DNA (10 minutes in a microcentrifuge, 40 minutes in a swinging-bucket centrifuge). The pellet was then rinsed with 70% ethanol and then left to air dry before being resuspended in 200 ul nuclease-free water.

For higher throughput 96-well plate DNA extractions, we extracted DNA from three to four week old colonies using a filter-plate prep as described in Strable et al[5].

For genotyping PCRs, 1-2 ul of genomic DNA was used to PCR a small (200-300 bp) region flanking the gRNA target site. A subset of PCR products were checked on a gel, and PCR products were purified using ExoSap-IT (https://www.thermofisher.com/order/catalog/product/78205.10.ML#/78205.10.ML) PCR cleanup reagent before being sent for Sangar sequencing. Large sets of Sangar sequencing results were converted into FASTA format and aligned with the wild type sequence using MUSCLE[6].

#### Staining and Imaging

Slides were prepared by extruding a ring of vacuum grease through a syringe to onto glass slides. These rings retained a pool of staining solution and prevented gametophores from being crushed by the coverslip. The area within the vacuum grease was filled with 5 µg/m Propidium Iodide (PI). Tufts of moss grown on BCD were flooded with water and vigorously tapped to remove as many air bubbles as possible. Gametophores were then dissected from the periphery of tufts and placed into the PI solution on the slides. Moss was allowed to stain for at least 15 minutes in the PI solution before imaging. Imaging was conducted using a Zeiss 710 laser scanning confocal microscope. We used a 514 nm laser for excitation and collected an emission spectrum from 566 to 650 nM. Images were captured using either a 20x water immersion NA 1.0 or 10x air lens.

#### Ectopic Apical Cell Quantification

Areas of stem visible on maximum intensity projections (not including leaves) were manually measured using FIJI[7]. Ectopic apical cells were identified by eye as triangular cells sometimes surrounded by short, curled hair cells. Apical cells were distinguished from illusory tetrahedral cells seen in maximum intensity projections by manually looking through z-stacks. Regions of ectopic growth and leaf production were also counted as ectopic stem cells.

#### Statistical analysis

Statistical analysis was performed using the R statistical programming language. Genotypes were coded as two loci (*CLV1* and *RPK2*) for the purposes of our analysis, since *clv1a* and *clv1b* single mutants were not analyzed separately. The effects of mutating CLV1 and RPK2 and of growth on 10nM and 100nM BAP as well as the interactions between these three factors (the two loci and the media) were modeled using a Poisson general linear model that included stem area as an offset (model: glm(total_ectopic ~ clv1L +

rpk2L + exock + clv1L:rpk2L + clv1L:exock + rpk2L:exock + (log(area)), family = 'poisson'). BAP concentrations were treated as a continuous variable with levels of zero, ten, and one hundred.

Poisson coefficients are akin to Beta values reported by linear regressions, in that they are proportional to the expected change in the dependent variable given the change in independent variable associated with the coefficient. In the case of a Poisson coefficient, the exponentiation of the coefficient tells you the predicted effect due to the change in factor level. For instance, with a Poisson coefficient of 0.64, the estimated change in apical cell number due to the *clv1* mutation is ~1.9 ( = e^0.64). It is important to note that our models make use of both categorical and continuous variables, which makes the coefficients appear deceptively different in magnitude. For example, the coefficient associated with exogenous cytokinin is small because cytokinin is coded as a continuous variable. The coefficient is 0.013 and it's exponent is 1.013, which appears much lower than the expected change due to *clv1* of 1.9. However, the cytokinin coefficient of 1.013 shows the predicted change per unit cytokinin. The predicted change for 10nM BAP is the exponentiation of 10*the coefficient, so e^(10*.013) = 1.14. Going on to predict the change for 100nM bap is e^(1.3) = 3.67. Finally, these numbers represent the fold change from the 'intercept' value also reported by the model.

#### gRNA and primers

| | | |
|---|---|---|
| oJCm278 | GAGTTAGGGGAGATGACGCG | rpk2_gRNA target locus genotyping |
| oJCm279 | CTTGGAGGACTCACCAACCC | rpk2_gRNA target locus genotyping |
| oJCm379 | cacctaaacggctcaattcc | PpClv1aE4 genotyping |
| oJCm380 | tgatgatctccgatggtatgg | PpClv1aE4 genotyping |
| oJCm181 | tggagagacgcaacttccat | CLL1b_exon1_sample - sgRNAs1 and 2 |
| oJCm182 | ttaagacgccccaaatcagc | CLL1b_exon1_sample - sgRNAs1 and 2 |
| oJCm175 | gcttcGAGCTCGAATTCAGA | PpU3_promoter_fwd for Sanger sequencing of gRNA plasmids |
| oJCm176 | ggtcGACGAGCTCAAAAAAAG | sgRNA_scaffold_reverse for Sanger sequencing of gRNA plasmids |
| oJCm208 | GAGCTCGAATTCGTCCATTGA | U6 promoter fwd pcr primer for Sanger sequencing of gRNA plasmids |

| **sgRNA Oligo for synthesis** | **Sequence** | **Target** | **Inserted into Vector** |
|---|---|---|---|
| sgJTC5 | GGCAgacagtgcccgaggctctct | CLL1a_exon4_cds | U3_BSAI-sgRNA |
| sgJTC6 | AAACagagagcctcgggcactgtc | CLL1a_exon4_cds* | U3_BSAI-sgRNA |

| | | | |
|---|---|---|---|
| sgJTC9 | GGCagaagtgcgagaccctcttc | CLL1b_exon1_cds_sgRNA1 | U3_BSAI-sgRNA |
| sgJTC10 | AAACgaagagggtctcgcacttc | CLL1b_exon1_cds_sgRNA1* | U3_BSAI-sgRNA |
| sgJTC105 | catGGGTTTGAGCGACGATGGCC | PpRPK2cds | U6_sgRNA |
| sgJTC106 | aaacGGCCATCGTCGCTCAAACCC | PpRPK2cds | U6_sgRNA |

#### Genes referenced in this study

| **Full Gene Name** | **Alias** | **Version 1.6** | **Version 3** |
|---|---|---|---|
| *CLAVATA1a* | *CLV1a* | Pp1s14_447V6 | Pp3c6_21940 |
| *CLAVATA1b* | *CLV1B* | Pp1s5_68V6 | Pp3c13_13360 |
| *RECEPTOR-LIKE PROTEIN KINASE 2* | *RPK2* | Pp1s311_57V6 | Pp3c7_5570 |
| *CYTOKININ HISTIDINE KINASE 1* | *CHK1* | Pp1s50_141V6 | Pp3c25_8540 |
| *CYTOKININ HISTIDINE KINASE 2* | *CHK2* | Pp1s194_72V6 | Pp3c16_7610 |
| *CYTOKININ HISTIDINE KINASE 3* | *CHK3* | Pp1s252_49V6 | Pp3c6_7030 |

### Dynamical Model Methods

Each of the models described here simulated the accumulation of gene products through time, simultaneously modeling transcription and translation. The equations are modified from Gordon et al. 2009, where the authors use similar systems of differential equations to test predictions about CLV43, CLV1, WUS, and cytokinin interactions[8].

#### Summary of workflow

1) Run the model and confirm that it converges to a steady-state value within the allotted steps.
2) Simulate each mutant genotype at each cytokinin level of interest with the initial parameters to generate a starting fit score.
3) Begin the optimizer: randomly mutate parameters, compare fit score, repeat
4) After 300 runs of the optimizer, extract parameters from optimal run and plot simulations vs. empirical data

#### Model Variables

Dynamical models were systems of Ordinary Differential Equations (ODEs) based on equations used to model cytokinin-CLV dynamics in Gordon et al[8]. The set of differential equations that constitutes a model describes the change in a set of interrelated variables through time. The variables used in this work are summarized here:

| Variable | Describes |
|---|---|
| x | Cytokinin-response pathway that induces stem cell formation |
| y | Cytokinin-independent pathway inducing stem cell formation |
| z | Cytokinin feedforward control of stem cell formation |
| init | Level of stem cell initiation |
| clv | Strength of CLV1 signaling. This is a static, non-dynamical parameter |
| rpk2 | Strength of RPK2 signaling. This is a static, non-dynamical parameter |
| cyt | Strength of cytokinin signaling. Set to 0 for *chk*, 1 for mock-treated wt, and to 10 and 100 for cytokinin treatments |

It is important to note that these variables are not meant to exactly reflect the level of a protein, but more the presence/absence and strength of the signaling pathway.

#### Model parameters

Each equation in the model describes how one of the above variables changes over time. The change over time is proportional to the current value of the parameters and other variables in the model. Each of the other variables in an equation is associated with a proportionality constant that describes how that variable affects the accumulation rate described by that equation. Additionally, a differential equation might include a term to describe accumulation independent of the other variables as well as degradation rates. These constants were assigned to the following categories:

- p = production; describes basal accumulation rates
- d = degradation; describes degradation rates
- k = interaction coefficient/proportionality constant

Each model also used a set of initial conditions, which we named 'base' values, and a time vector that we named times0. Models were run for 3000 time points distributed over 300 'seconds'. Finally, cytokinin was coded as a static parameter and altered in the following ways to simulate different conditions from our experimental datasets:

| Cytokinin value | Simulates the condition |
|---|---|
| 0 | *chk* triple mutant |
| 1* | growth on minimal media (BCD) with wild type *CHK* genes |
| 10 | 10 nM BAP |
| 100 | 100 nM BAP |

* As '1' here is somewhat arbitrary, we also tried values of 0.5 and 0.75 in its stead, to no significant change to the model outputs (not shown).

#### Running a model

Each model was solved using the LSODA solver for Ordinary Differential Equations (ODEs) and the R statistical programming language Version 4.0.2[9,10]. Models were confirmed to converge to steady state values before and after each run of the optimizer, as determined by each variable reaching a plateau by the end of the modeled time period. All plotting used the ggplot2 package[11]. Models were run for 2000 or 3000 time points (steps) distributed over 200 300 'seconds', as depicted by the sample model run below. Variables change through time and converge at steady state values. The final values at time 200 or 300 (more steps were given to models that took longer to converge) were taken and stored as the output of the model

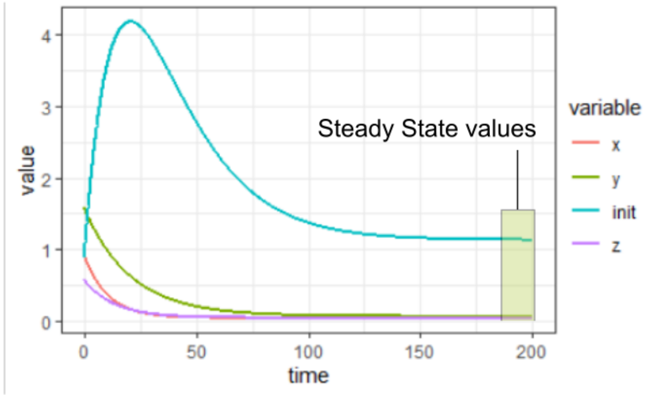


Variables change through time and converge at steady state values (above figure). The final values, at time 300 (the 3,000th time point used for integration) were taken and stored as the output of the model.

#### Normalization and fit to the empirical data

Comparing the simulated data to the empirical data required that the two datasets be normalized to a unified scale. To achieve this, the empirical data set was normalized to the ectopic stem cell per area value of wild type moss grown on minimal media. For the model, 'area' was not considered, and the modeled stem cell initiation values (termed init) were also normalized to the modeled stem cell values of wild type on minimal media (cytokinin = 1). This allowed us to compare the trends in the data, for instance if *clv1* mutants on minimal media made four times as many stem cells per area as wild type on minimal media, we assigned a value of four to this condition in the dataset. The model would then attempt to generate a value for *clv1* at the minimal media cytokinin input parameter that was four times higher than wild type at the same cytokinin level.

Each simulated value was compared to its corresponding empirical data value to generate a score. These scores were used to penalize a model with a given set of parameters; higher scores were worse than lower scores. The score was intended to accomplish the following:

1) Equally penalize simulated values that overshot or undershot the data

2) Weigh all datapoints equally, regardless of magnitude. To do this, the score had to **minimize fold changes between the simulated and empirical data**. Otherwise, a change from 1->2 would be penalized less than a change from 10 to 14, despite the former constituting a much larger relative change.
3) Penalize larger deviations from the data more severely than smaller ones. Otherwise, a model might be 'optimized' to have good fits to some data points but terrible fits to others. Since the intention of the model is to capture the trends in the data across all conditions, such a scenario was unacceptable.

We used the log of the fold change between the simulated ($m_i$) and empirical data ($d_i$) to accomplish the above aims one and two, and then squared to accomplish aim 3. The sum of these penalty scores at each data point ($P_i$) then yielded the total fit score *F*:

$$P_i = \ln\left(\frac{m_i}{d_i}\right)$$

$$F = \sum {P_i}^2$$

#### Mutating parameter values

Each model was run initially with semi-arbitrary parameters that allowed the model to converge within the given time frame. When comparing models, each model used similar starting parameters before optimization.

After each run, new model parameters *k, p*, *d*, and *base* were randomly selected from a normal distribution based around the previous parameter value and the model was run again. The Fit Score *F* for the new model was compared with the previous *F*. If the new *F* proved lower than the previous, then the new model parameters were saved and mutated again for the next run. If instead the new *F* was not lower than the previous, the original parameter set was randomly mutated again.

To mutate the parameters, a vector of values following a normal distribution and centered on 1 was generated. The vector was the same length as the number of parameters to be mutated. The standard deviation of the distribution was 0.1.

#### Simulating mutant genotypes

To simulate mutant genotypes in models 1-7, CLV1 and RPK2 were non-dynamical, we set CLV1 or RPK2 to and their synthesis parameters to 0. *chk* mutants were simulated by setting the 'cytokinin' input parameter to 0.

#### Model 1-5

Models 1-5 consist of the following equations. Edges in the network (such as RPK2 inhibition of y) were changed by setting corresponding *k* values (in this case, *k*[5], to 0).

$$\frac{dx}{dt} = \frac{p_1 + cyt * k_1}{1 + p_1 + cyt * k_1 + k_2 * clv + k_3 * rpk2} - d_1 * x$$

$$\frac{dy}{dt} = \frac{p_2}{1 + p_2 + k_4 * clv + k_5 * rpk2} - y * d_2$$

$$\frac{dinit}{dt} = \frac{p_3 + x * k_6 + y * k_7}{1 + p_3 + x * k_6 + y * k_7} - init * d_3$$

#### Model 6 and 7

Models 6 and 7 are similar in topology with the inclusion of the variable *z* downstream of cytokinin. In Model 6, *z* is inhibited by cytokinin, and induces *init*. In model 7, *z* is induced by cytokinin, and inhibits *init*. Two versions of each model were run: one with CLV1 inhibiting *x* and RPK2 inhibiting *y*, and one RPK2 inhibits *x* and CLV1 inhibits *y*.

#### Model 6

$$\frac{dx}{dt} = \frac{p_1 + cyt * k_1}{1 + p_1 + cyt * k_1 + clv * k_2 + rpk2 * k_3 + z * k_9} - x * d_1$$

$$\frac{\mathrm{dy}}{\mathrm{dt}} = \frac{p_2}{1 + p_2 + clv * k_4 + rpk2 * k_5} - \mathrm{y} * \mathrm{d}_2$$

$$\frac{dz}{dt} = \frac{p_4}{1 + p_4 + cyt * k_8} - z * d_4$$

#### Model 7

$$\frac{dx}{dt} = \frac{p_1 + cyt * k_1}{1 + p_1 + cyt * k_1 + clv * k_2 + rpk2 * k_3 + z * k_9} - x * d_1$$

$$\frac{dz}{dt} = \frac{p_4 + cyt * k_8 * cyt}{1 + p_4 + cyt * k_8} - z * d_4$$

$$\frac{dy}{dt} = \frac{p_2}{1 + p_2 + clv * k_4 + rpk2 * k_5} - y * d_2$$

$$\frac{dinit}{dt} = \frac{p_3 + x * k_6 + y * k_7}{1 + p_3 + x * k_6 + y * k_7 + z * k_9} - init * d_3$$

#### List of parameters, starting values, and finishing values for a sample run

##### Model 1 (Figure 4 A)

| Parameter | Description | Starting Value | Finishing Value |
|---|---|---|---|
| $k_1$ | cytokinin ↑ x | 1 | 1.500644 |
| $k_2$ | clv ↓ x | 1 | 0.525365 |
| $k_3$ | rpk2 ↓ x | 1 | 20.8025 |
| $k_4$ | clv ↓ y | 1 | 4.652118 |
| $k_5$ | rpk2 ↓ y | 1 | 0.190285 |
| $k_6$ | x ↑ init | 0.1 | 0.019766 |
| $k_7$ | y ↑ init | 0.1 | 0.319337 |
| $p_1$ | basal x synthesis | .01 | 0.00365 |
| $p_2$ | basal y synthesis | 0.01 | 0.02994 |
| $p_3$ | basal init synthesis | 0.01 | 0.008979 |
| $d_1$ | x degradation | 0.05 | 0.035265 |
| $d_2$ | y degradation | 0.05 | 0.035136 |
| $d_3$ | init degradation | 0.05 | 0.02294 |
| $base_1$ | initial x | 1 | 1.165125 |

| | | | |
|---|---|---|---|
| $base_2$ | initial y | 1 | 0.84229 |
| $base_3$ | initial init | 1 | 0.402657 |

#### Model 2 (Figure 4 B)

| Parameter | Description | Starting Value | Finishing Value |
|---|---|---|---|
| $k_1$ | cytokinin ↑ x | 1 | 0.304867 |
| $k_2$ | clv ↓ x | 1 | 5.506337 |
| $k_3$ | rpk2 ↓ x | 0 | 0 |
| $k_4$ | clv ↓ y | 0 | 0 |
| $k_5$ | rpk2 ↓ y | 1 | 3.955426 |
| $k_6$ | x ↑ init | 0.1 | 0.048441 |
| $k_7$ | y ↑ init | 0.1 | 0.209578 |
| $p_1$ | basal x synthesis | 0.1 | 0.025085 |
| $p_2$ | basal y synthesis | 0.1 | 0.0416 |
| $p_3$ | basal init synthesis | .01 | 0.003168 |
| $d_1$ | x degradation | 0.01 | 0.047735 |
| $d_2$ | y degradation | 0.01 | 0.017703 |
| $d_3$ | init degradation | 0.05 | 0.041424 |
| $base_1$ | initial x | 0.05 | 0.452641 |
| $base_2$ | initial y | 0.05 | 0.333642 |
| $base_3$ | initial init | 1 | 0.351427 |

#### Model 3 (Figure 4 C)

| Parameter | Description | Starting Value | Finishing Value |
|---|---|---|---|
| $k_1$ | cytokinin ↑ x | 1 | 0.655062 |
| $k_2$ | clv ↓ x | 0 | 0 |
| $k_3$ | rpk2 ↓ x | 1 | 7.480788 |
| $k_4$ | clv ↓ y | 1 | 2.17663 |
| $k_5$ | rpk2 ↓ y | 0 | 0 |
| $k_6$ | x ↑ init | 0.1 | 0.055229 |
| $k_7$ | y ↑ init | 0.1 | 0.27232 |
| $p_1$ | basal x synthesis | 0.1 | 0.014258 |
| $p_2$ | basal y synthesis | 0.1 | 0.015025 |
| $p_3$ | basal init synthesis | .01 | 0.002324 |
| $d_1$ | x degradation | 0.01 | 0.159216 |
| $d_2$ | y degradation | 0.01 | 0.03027 |
| $d_3$ | init degradation | 0.05 | 0.04435 |
| $base_1$ | initial x | 0.05 | 1.239331 |
| $base_2$ | initial y | 0.05 | 0.899113 |
| $base_3$ | initial init | 1 | 0.540307 |

#### Model 4 (Figure 4 D)

| Parameter | Description | Starting Value | Finishing Value |
|---|---|---|---|
| $k_1$ | cytokinin ↑ x | 1 | 0.841983 |

| | | | |
|---|---|---|---|
| $k_2$ | clv ↓ x | 1 | 2.50779 |
| $k_3$ | rpk2 ↓ x | 1 | 9.162987 |
| $k_4$ | clv ↓ y | 0 | 0 |
| $k_5$ | rpk2 ↓ y | 0 | 0 |
| $k_6$ | x ↑ init | 0.1 | 0.218686 |
| $k_7$ | y ↑ init | 0.1 | 0.070918 |
| $p_1$ | basal x synthesis | 0.1 | 0.009767 |
| $p_2$ | basal y synthesis | 0.1 | 0.004626 |
| $p_3$ | basal init synthesis | .01 | 0.003158 |
| $d_1$ | x degradation | 0.01 | 0.057663 |
| $d_2$ | y degradation | 0.01 | 0.106677 |
| $d_3$ | init degradation | 0.05 | 0.019892 |
| $base_1$ | initial x | 0.05 | 0.072355 |
| $base_2$ | initial y | 0.05 | 0.866898 |
| $base_3$ | initial init | 1 | 0.740589 |

#### Model 5 (Figure 4 E)

| Parameter | Description | Starting Value | Finishing Value |
|---|---|---|---|
| $k_1$ | cytokinin ↑ x | 1 | 0.035789 |
| $k_2$ | clv ↓ x | 0 | 0 |
| $k_3$ | rpk2 ↓ x | 0 | 0 |
| $k_4$ | clv ↓ y | 1 | 0.296071 |
| $k_5$ | rpk2 ↓ y | 1 | 6.576975 |
| $k_6$ | x ↑ init | 0.1 | 0.027982 |
| $k_7$ | y ↑ init | 0.1 | 0.359106 |
| $p_1$ | basal x synthesis | 0.1 | 0.004176 |
| $p_2$ | basal y synthesis | 0.1 | 0.03289 |
| $p_3$ | basal init synthesis | .01 | 0.000394 |
| $d_1$ | x degradation | 0.01 | 0.045523 |
| $d_2$ | y degradation | 0.01 | 0.030657 |
| $d_3$ | init degradation | 0.05 | 0.045014 |
| $base_1$ | initial x | 0.05 | 0.551109 |
| $base_2$ | initial y | 0.05 | 0.300094 |
| $base_3$ | initial init | 1 | 0.702136 |

### Models 6 and 7

#### Model 6 (CLV inhibits x, Figure 5)

| Parameter | Description | Starting Value | Finishing Value |
|---|---|---|---|
| $k_1$ | cytokinin ↑ x | 1 | 0.435963 |
| $k_2$ | clv ↓ x | 1 | 5.048187 |
| $k_3$ | rpk2 ↓ x | 0 | 0 |
| $k_4$ | clv ↓ y | 0 | 0 |
| $k_5$ | rpk2 ↓ y | 1 | 8.082593 |
| $k_6$ | x ↑ init | 0.1 | 0.017249 |
| $k_7$ | y ↑ init | 0.1 | 0.169828 |
| $k_8$ | cytokinin ↓ z | 0.5 | 8.181178 |
| $k_9$ | z ↑ init | 0.5 | 0.768938 |
| $p_1$ | basal x synthesis | 0.01 | 0.004266 |
| $p_2$ | basal y synthesis | 0.03 | 0.023539 |
| $p_3$ | basal init synthesis | .01 | 0.004809 |
| $p_4$ | basal z synthesis | 0.01 | 0.008215 |
| $d_1$ | x degradation | 0.05 | 0.04629 |
| $d_2$ | y degradation | 0.05 | 0.027294 |
| $d_3$ | init degradation | 0.05 | 0.032801 |
| $d_4$ | z degradation | 0.05 | 0.063203 |
| $base_1$ | initial x | 1 | 1.927968 |
| $base_2$ | initial y | 1 | 0.232216 |
| $base_3$ | initial init | 1 | 0.379397 |
| $base_4$ | initial z | 1 | 0.656615 |

#### Model 6 (RPK2 inhibits X)

| Parameter | Description | Starting Value | Finishing Value |
|---|---|---|---|
| $k_1$ | cytokinin ↑ x | 1 | 0.380677 |
| $k_2$ | clv ↓ x | 0 | 0 |
| $k_3$ | rpk2 ↓ x | 1 | 6.363934 |
| $k_4$ | clv ↓ y | 1 | 2.772667 |
| $k_5$ | rpk2 ↓ y | 0 | 0 |
| $k_6$ | x ↑ init | 0.1 | 0.053368 |
| $k_7$ | y ↑ init | 0.1 | 0.205299 |
| $k_8$ | cytokinin ↓ z | 0.5 | 12.45164 |
| $k_9$ | z ↑ init | 0.5 | 0.854566 |
| $p_1$ | basal x synthesis | 0.01 | 0.059793 |
| $p_2$ | basal y synthesis | 0.03 | 0.032342 |
| $p_3$ | basal init synthesis | .01 | 0.002267 |
| $p_4$ | basal z synthesis | 0.01 | 0.008643 |
| $d_1$ | x degradation | 0.05 | 0.097205 |
| $d_2$ | y degradation | 0.05 | 0.048109 |
| $d_3$ | init degradation | 0.05 | 0.052553 |
| $d_4$ | z degradation | 0.05 | 0.074966 |
| $base_1$ | initial x | 1 | 0.933395 |
| $base_2$ | initial y | 1 | 1.584061 |
| $base_3$ | initial init | 1 | 0.876901 |
| $base_4$ | initial z | 1 | 0.579399 |

#### Model 7 (CLV inhibits X, Figure 5)

| Parameter | Description | Starting Value | Finishing Value |
|---|---|---|---|
| $k_1$ | cytokinin ↑ x | 1 | 0.863826 |
| $k_2$ | clv ↓ x | 1 | 19.77114 |
| $k_3$ | rpk2 ↓ x | 0 | 0 |
| $k_4$ | clv ↓ y | 0 | 0 |
| $k_5$ | rpk2 ↓ y | 1 | 1.999533 |
| $k_6$ | x ↑ init | 0.1 | 0.040456 |
| $k_7$ | y ↑ init | 0.1 | 0.173769 |
| $k_8$ | cytokinin ↑ z | 0.5 | 1.685863 |
| $k_9$ | z ↓ init | 0.5 | 0.311607 |
| $p_1$ | basal x synthesis | 0.01 | 0.028438 |
| $p_2$ | basal y synthesis | 0.03 | 0.056778 |
| $p_3$ | basal init synthesis | .01 | 0.004693 |
| $p_4$ | basal z synthesis | 0.01 | 0.005662 |
| $d_1$ | x degradation | 0.05 | 0.014775 |
| $d_2$ | y degradation | 0.05 | 0.018722 |
| $d_3$ | init degradation | 0.05 | 0.038537 |
| $d_4$ | z degradation | 0.05 | 0.118948 |
| $base_1$ | initial x | 1 | 0.654166 |
| $base_2$ | initial y | 1 | 0.251628 |

| | | | |
|---|---|---|---|
| $base_3$ | initial init | 1 | 2.202736 |
| $base_4$ | initial z | 1 | 0.992124 |

##### Model 7 (RPK2 inhibits X)

| Parameter | Description | Starting Value | Finishing Value |
|---|---|---|---|
| $k_1$ | cytokinin ↑ x | 1 | 1.463869 |
| $k_2$ | clv ↓ x | 0 | 0 |
| $k_3$ | rpk2 ↓ x | 1 | 19.17415 |
| $k_4$ | clv ↓ y | 1 | 1.013134 |
| $k_5$ | rpk2 ↓ y | 0 | 0 |
| $k_6$ | x ↑ init | 0.1 | 0.027886 |
| $k_7$ | y ↑ init | 0.1 | 0.087528 |
| $k_8$ | cytokinin ↑ z | 0.5 | 2.157959 |
| $k_9$ | z ↓ init | 0.5 | 0.190479 |
| $p_1$ | basal x synthesis | 0.01 | 0.039586 |
| $p_2$ | basal y synthesis | 0.03 | 0.061077 |
| $p_3$ | basal init synthesis | .01 | 0.00435 |
| $p_4$ | basal z synthesis | 0.01 | 0.012878 |
| $d_1$ | x degradation | 0.05 | 0.012762 |
| $d_2$ | y degradation | 0.05 | 0.024692 |
| $d_3$ | init degradation | 0.05 | 0.055329 |
| $d_4$ | z degradation | 0.05 | 0.042331 |
| $base_1$ | initial x | 1 | 0.432841 |
| $base_2$ | initial y | 1 | 1.570673 |
| $base_3$ | initial init | 1 | 0.574236 |
| $base_4$ | initial z | 1 | 0.270798 |

**Media**
**General Moss Stock Solutions (100x)**
**Stock solution B:**
MgSO4.7H2O (magnesium sulphate 7-hydrate) 2.5 g
(or 1.2 g of anhydrous MgSO4)
dH2O to 100 ml

**Stock solution C:**
KH2PO4 (potassium phosphate) 2.5 g
dH2O to 50 ml
Adjust pH to 6.5 with minimal volume of 4 M KOH. Then make up to 100 ml with additional dH2O.

**Stock solution D:**
KNO3 (potassium nitrate) 10.1 g
FeSO4.7H2O (iron sulphate 7-hydrate) 0.125 g
dH2O to 100 ml

**0.5 M di-ammonium (+) tartrate:**
9.2 g in 100 ml dH2O

**20x trace element solution**:
H3BO3 (boric acid) 614 mg
AlK(SO4)2.12H2O (aluminium potassium
sulphate 12-hydrate) 55 mg
CuSO4.5H2O (cupric sulphate 5-hydrate) 55 mg
KBr (potassium bromide) 28 mg
LiCl (lithium chloride) 28 mg

$MnCl_2.4H_2O$ (manganese chloride 4-hydrate) 389 mg
$CoCl_2.6H_2O$ (cobalt chloride) 55 mg
$ZnSO_4.7H_2O$ (zinc sulphate 7-hydrate) 55 mg
KI (potassium iodide) 28 mg
$SnCl_2.2H_2O$ 28 mg
$dH_2O$ to 50 ml

**0.5 M $CaCl_2$:**
3.67g $CaCl_2$ in 50 ml dH20

**Solutions for transformation**
**8.5% Mannitol** (Sigma M1902-1KG)
85 g in 1 liter

**Driselase** for Protoplasting (200ml)
4 g Driselase into 200 ml 8.5% Mannitol.
Gently stir for 30 minutes at room temperature.
Keep at 4°C for 30 minutes.
Stir 5 minutes at room temperature.
Spin at 2,500g for 10 minutes in 50 ml Falcon Tubes.
Filter sterilize with 0.22 µm filter.
Aliquot 10 ml into 15 ml Falcon Tubes.

**3M Solution (50ml, store at 4°C)**

| | |
|---|---|
| 4.55 g | Mannitol |
| 750 µl | 1M $MgCl_2·6H_2O$ |
| 5 ml | 1% MES pH 5.6 |
| | $H_2O$ to **50** ml |

Vortex well
Filter sterilize with 0.22 µm filter
Store at 4°C

**PEG Solution for Transformation (use right away or store at -20°C)**

1) **Prepare Man/$Ca(NO_3)_2$ Solution**

| | |
|---|---|
| 9 ml | 8.5% Mannitol |
| 1 ml | 1M $Ca(NO_3)_2·4H_2O$ |
| 100 µl | 1M Tris pH 8.0 |

Vortex well. Filter sterilize with 0.22 µm filter.

2) **Prepare PEG**

4 g PEG 8000 (Sigma P-2139) into 50 ml Falcon Tube or in a glass vial (preferable, as plastic tube can melt).

Melt in microwave, watching carefully.

**Combine**

In sterile hood add 1 and 2 together (mannitol/$Ca^{2+}$ and PEG). Vortex well. Make sure all is well mixed. Solution can be used after 2 hours or can be stored at –20°C for long-term storage.

**PRMB**

PRMB is BCDAT media with the following:
6% (w/v) Mannitol
0.55% Agar
After autoclaving: add 1ml 500 mM $CaCl_2$ per 50 ml of media.

**PRMT**
PRMT is **BCDAT** media with the following added:
6% (w/v) Mannitol
0.3% agar
After microwaving: add 1ml 500 mM $CaCl_2$ per 50 ml of media.

10. Team, R. C. R: A Language and Environment for Statistical Computing. *R Foundation for Statistical Computing* (2016).

11. Hadley, W. *et al.* *ggplot2 - Elegant Graphics for Data Analysis*. *ggplot2: Elegant Graphics for Data Analysis* (Springer-Verlag New York, 2016).
